# Selection of forage oat genotypes through GGE Biplot and BLUP

**DOI:** 10.1101/2020.03.10.986422

**Authors:** Franklin Santos, y Félix Marza

## Abstract

In Bolivia, there is a low predominance of forage oat productivity. Therefore, it was proposed to select more productive and stable genotypes through statistical methods of GGE Biplot and BLUP. The research was conducted in three environments in Bolivia and six commercial varieties of forage oats were evaluated; three of them correspond to INIA Peru and the rest of Bolivia. Data were analyzed through GGE Biplot and BLUP (Best Linear Unbiased Prediction) and an average yield of 10.29 ±3.51 t ha^−1^ of dry matter was obtained. BLUP accumulated greater variance than GGE Biplot in the first two components. In terms of productivity and stability values, both models have the same selection trend. Thus, Tayco and Texas were selected for their outstanding characteristic in dry matter yield and phenotypic stability.

## Introduction

Bolivia produced 11,363.0 tons of forage oats in 2018, with a yield of 2,464.0 kg ha^−1^ (INE & MDYT, 2019). These data reflect low productivity that prevails in our country compared to neighboring countries in the region. In Bolivian highlands, approximately 95% of biomass production is stored in haylage, for supply in winter season, since adequate and timely food availability determine the success and prosperity of livestock (Bilal, Ayub, Tariq, Tahir, & Nadeem, 2017) and insufficient forage supply could reduce meat and milk productivity (Ahmad et al., 2014; Rana et al., 2014). To increase production, plant breeding is a key mechanism to identify genotypes of high productivity and resilience to biotic and abiotic adverse factors (Atlin, Cairns, & Das, 2017). This process requires a multi-environmental tests to select or recommend genotypes that combine high yields and stability in time and space (Tiago Olivoto, Lúcio, da Silva, Sari, & Diel, 2019; Smith & Cullis, 2018). This requires applying statistical models with better prediction skills.

In multi-environmental analyzes there are statistical models to study adaptability and stability of cultivars, these methodologies are based on analysis of variance, nonparametric regressions, multivariate analysis, and mixed models. Thus, GGE Biplot and REML/BLUP (Best Linear Unbiased Prediction) stand out for practicality in their interpretation of results and precision in selection of genotypes (T Olivoto et al., 2017; A. d. Santos, Amaral Júnior, Kurosawa, Gerhardt, & Fritsche Neto, 2017). GGE biplot graphics are constructed from first and second main components, that represent the genotype ratio and G×E interaction, allowing accurate prediction of the average genotype yield in different environments and identifying stable genotypes (Yan & Tinker, 2006). On the other hand, Resende (2016) and P. Santos et al. (2019) state that BLUP is an optimal selection procedure that maximizes selective accuracy and allows the simultaneous use of several sources of information, addresses the imbalance, considers the genetic relationship between the plants evaluated and the coincidence between the selection and recombination units.

Mixed models have been widely used in plant breeding programs for their estimation procedures reduce unbalanced design skewness, non-additive features or could adjust scattered data from an investigation (Bandera-Fernández & Pérez-Pelea, 2018; Hu, 2015). Also, it can estimate important parameters of quantitative genetics from multi-environmental trials and select the best genotypes through productivity, adaptability, and stability.

AMMI (Main Additive Effect and Multiplicative Interaction) and BLUP are popular methods for analyzing trials in multi-environments; However, AMMI has deficiencies in accommodating a linear mixed-effect model (LMM) structure and BLUP needs to deal graphically with a random GEI structure (Tiago Olivoto, Lúcio, da Silva, Marchioro, et al., 2019). WAASB is the weighted average of absolute scores from decomposition of the singular value of the BLUP matrix of interaction effects (GEI) generated by an LMM proposed by Tiago Olivoto, Lúcio, da Silva, Marchioro, et al. (2019), in which, they mention that WAASB is an important statistical tool to identify highly productive and widely adapted genotypes. Therefore, through the GGE Biplot and BLUP methods, it was proposed to select more productive and stable genotypes of forage oats in three environmental conditions in Bolivia.

## Materials and methods

The research was carried out in three locations in Bolivia (Table 1). Cuyahuani is located in a north highlight of La Paz. In this area, the families are engaged in milk production; The Toralapa Innovation Center is located in the upper part of Cochabamba on the old Santa Cruz road, and the National Potato Innovation Center is located in the high valley of Cochabamba, the economic activity in both sectors is agriculture.

**Table 1.**
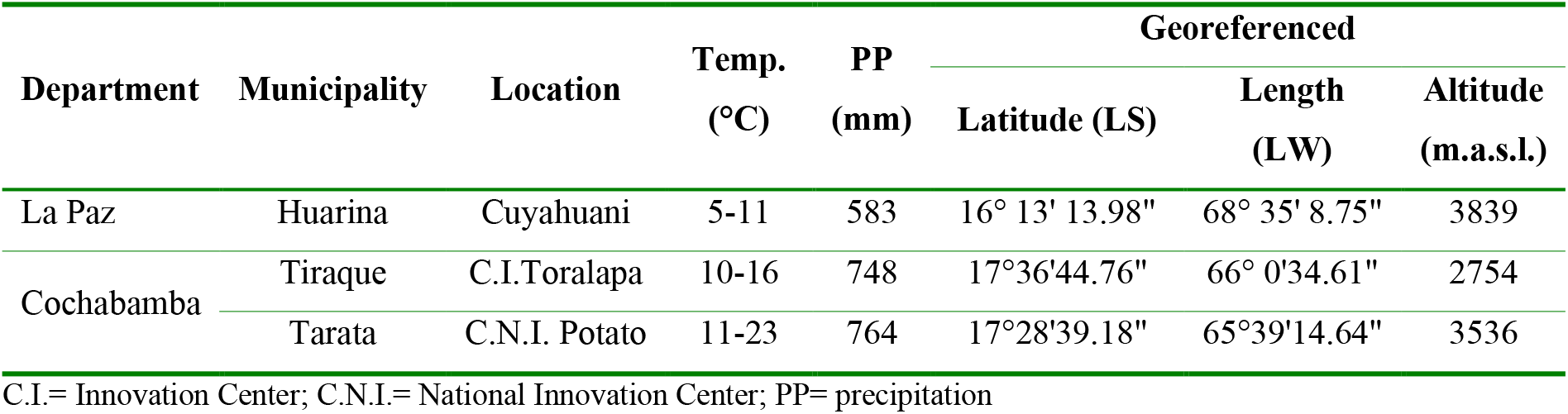
Location corresponding to the research works where they were implemented during the 2018-2019 agricultural campaign.

The statistical delineation of research in three environments was completely randomized blocks with 3 repetitions. The experimental plot was composed of five rows of 5 m and a distance of 0.25 m between rows. The sowing was carried out with a continuous jet at a density of 90 kg ha^−1^. The genetic material evaluated consisted of six varieties of forage oats, three of them were obtained from INIA Peru (INIA-Tayco, INIA-Villcanota, and INIA-Africana), the remaining three were acquired from the CIF-Violeta Seed Company, located in Cochabamba Department, Bolivia (Texas, Gaviota, and Aguila). The type of tillage in the trials was conventional, in a rotating soil. The crop was kept free of weeds, by manual control.

The harvest was carried out when each genotype reached 50% of the pasty grain and the sample was weighed to obtain green matter yield. The dry matter content in percentage (%) was determined from a sub-sample (500 g) of fresh fodder that was taken in a moisture-free kraft paper bag and dried in an oven at 105 °C until obtaining a weight constant minimum. The resulting value was multiplied by the fresh mass to calculate the dry matter yield of each respective experimental unit.

To obtain the results, a descriptive analysis of measures of central tendency and dispersion was carried out; skewness and kurtosis were also determined as a normality test. To test statistical differences (P <0.01 and 0.05), a combined analysis of variance was performed. On the other hand, for the analysis of multi-environmental trials, methodologies of GGE biplot (Genotype, Genotype × Environment) and BLUP (Best Linear Unbiased Prediction) were used, both models are described in detail by T Olivoto (2019), which is available at https://tiagoolivoto.github.io/metan/articles/vignettes_gge.html#the-gge-model and https://tiagoolivoto.github.io/metan/articles/vignettes_blup.html. The data were processed through the R Statistical Analysis System (R Core Team, 2019), for the multi-environment analyses also known as MET (Multi-Environment Trial), the METAN package by Tiago Olivoto (2019) was used.

## Results and Discussion

In Table 1, the average of plant length the genotypes studied was 112.16 ±19.87 cm. The leaves of the middle and upper third of the genotypes had an average length of 25.02 ±9.46 cm and a width of 1.62 ±0.39 cm. Within the biomass productivity measures, the national average green matter yield is 2.3 t ha^−1^ (INE & MDRYT, 2019); however, in this investigation, an average of 41.30 ±17.57 t ha^−1^ of GM and 10.29 ±3.51 t ha^−1^ of DM were obtained, which are higher than the national average. On the other hand, lower values (±1) of skewness and kurtosis indicate normality (Mishra et al., 2019), therefore, the research values are within these parameters, except leaf length.

**Table 1.**
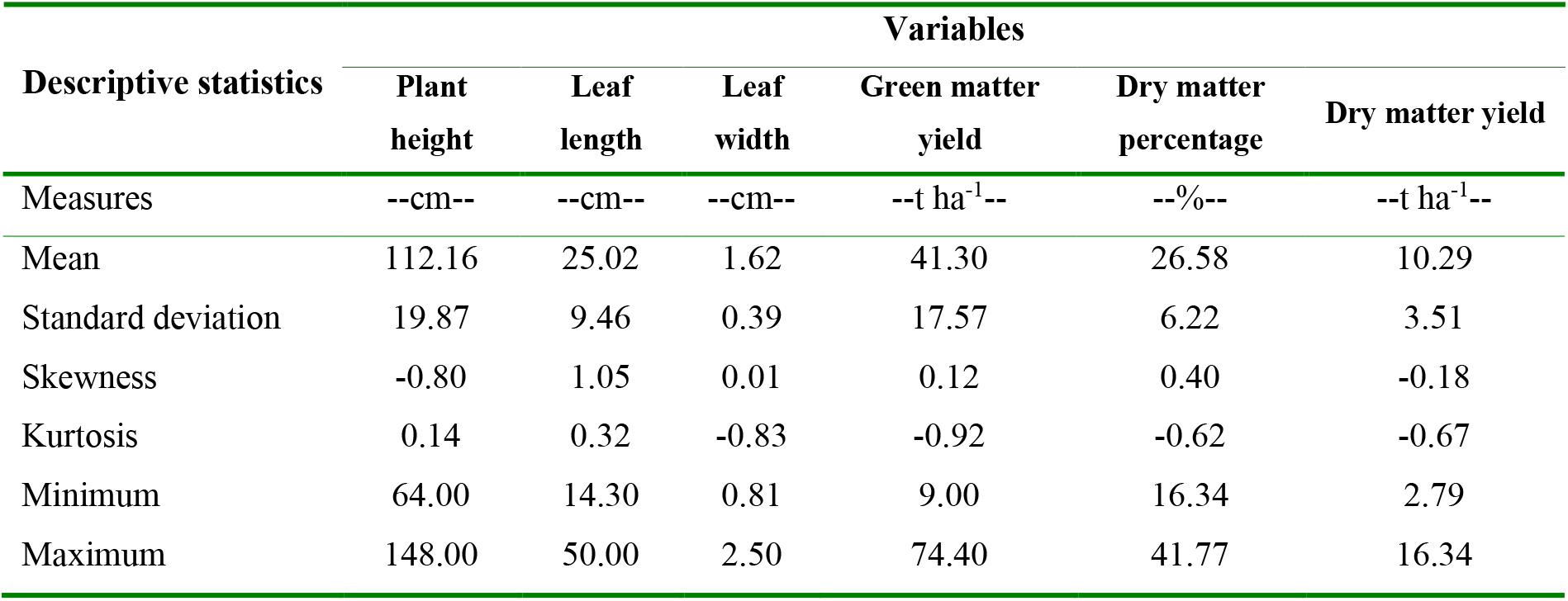
Descriptive statistics of six oats forage genotypes in localities of Huarina, Tarata and Toralapa Innovation Center, during the 2018-2019 agricultural campaign.

| Descriptive statistics | Variables |  |  |  |  |  |
| --- | --- | --- | --- | --- | --- | --- |
|  | Plant height | Leaf length | Leaf width | Green matter yield | Dry matter percentage | Dry matter yield |

| Measures | --cm-- | --cm-- | --cm-- | --t ha<sup>-1</sup>-- | --%-- | --t ha<sup>-1</sup>-- |
| --- | --- | --- | --- | --- | --- | --- |
| Mean | 112.16 | 25.02 | 1.62 | 41.30 | 26.58 | 10.29 |
| Standard deviation | 19.87 | 9.46 | 0.39 | 17.57 | 6.22 | 3.51 |
| Skewness | -0.80 | 1.05 | 0.01 | 0.12 | 0.40 | -0.18 |
| Kurtosis | 0.14 | 0.32 | -0.83 | -0.92 | -0.62 | -0.67 |
| Minimum | 64.00 | 14.30 | 0.81 | 9.00 | 16.34 | 2.79 |
| Maximum | 148.00 | 50.00 | 2.50 | 74.40 | 41.77 | 16.34 |

According to the combined analysis of variance (Table 2), the genotypes did not show differences (P> 0.05), except the leaf width (P<0.05); however, significant differences (P<0.01) were observed between environments referred to the variable plant height, leaf length, leaf width, green matter yield, and dry matter percentage. In this line, the non-significant interaction of the evaluated variables could indicate that the genotypes have similar behavior in the evaluated environments or it could be inferred that the environments do not have the sufficient capacity of discrimination. However, Fasahat, Rajabi, Mahmoudi, Noghabi, and Rad (2015) mentions that stability analyzes according to several principles can result in better identification of stable genotypes, even when there are no interactions between the parameters. Meanwhile, the coefficients of variation are within the acceptable ranges and the coefficients of determination showed their genetic expression above 50%.

**Table 2.**
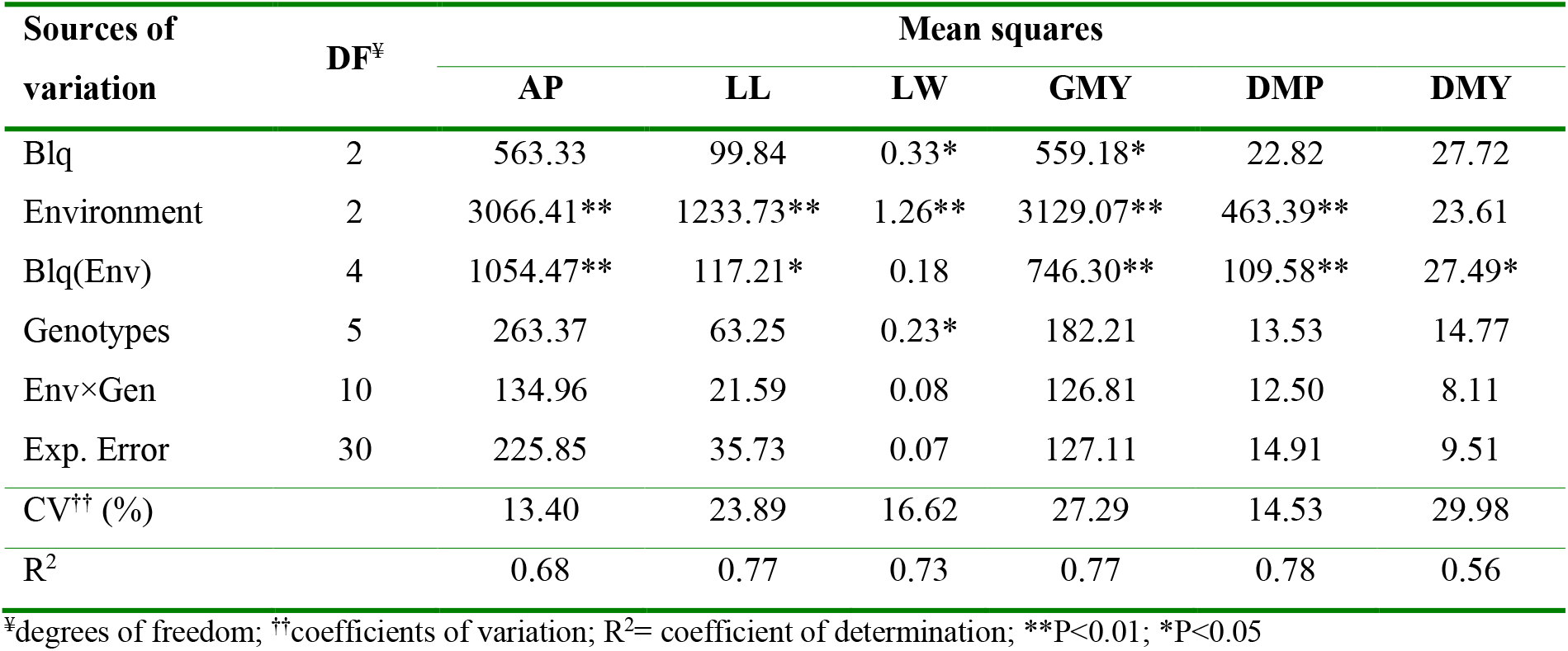
Analysis of variance for the variables PH (plant height), LL (leaf length), LW (leaf width), GMY (green matter yield), DMP (dry matter percentage) and DMY (dry matter yield).

| Sources of variation | DF<sup>y</sup> | Mean squares |  |  |  |  |  |
| --- | --- | --- | --- | --- | --- | --- | --- |
|  |  | AP | LL | LW | GMY | DMP | DMY |
| Blq | 2 | 563.33 | 99.84 | 0.33* | 559.18* | 22.82 | 27.72 |
| Environment | 2 | 3066.41** | 1233.73** | 1.26** | 3129.07** | 463.39** | 23.61 |
| Blq(Env) | 4 | 1054.47** | 117.21* | 0.18 | 746.30** | 109.58** | 27.49* |
| Genotypes | 5 | 263.37 | 63.25 | 0.23* | 182.21 | 13.53 | 14.77 |
| Env×Gen | 10 | 134.96 | 21.59 | 0.08 | 126.81 | 12.50 | 8.11 |
| Exp. Error | 30 | 225.85 | 35.73 | 0.07 | 127.11 | 14.91 | 9.51 |
| CV<sup>††</sup> (%) |  | 13.40 | 23.89 | 16.62 | 27.29 | 14.53 | 29.98 |
| R<sup>2</sup> |  | 0.68 | 0.77 | 0.73 | 0.77 | 0.78 | 0.56 |

<sup>y</sup>degrees of freedom; <sup>††</sup>coefficients of variation; R<sup>2</sup>= coefficient of determination; \*\* $P < 0.01$ ; \* $P < 0.05$

In GGE Biplot methodology, it was verified that the first two main components (PC1: 64.1% and PC2: 23.05%) accumulated a variance of 87.15%, derived from the decomposition of singular genotype values (G) + interaction effects (GxE). Yan and Kang (2002) mention that the first main component (PC1) shows the adaptability of genotypes that are highly correlated with performance, the second (PC2) indicates stability, that is, the genotypes closest to zero are the most stable. Accordingly, the Texas variety meets the stability criteria, as it is closer to the point of origin of the first component. On the other hand, Yan and Kang (2002) also indicate that the polygonal view of a biplot is the best way to visualize the patterns of interaction between genotypes and environments and interpret a biplot effectively. Therefore, the vertex genotypes in Figure 1b are Tayco, Aguila, Gaviota, Villcanota, and Africana, these varieties have a higher dry matter yield for the environments located in each sector.

**Figure 1.**
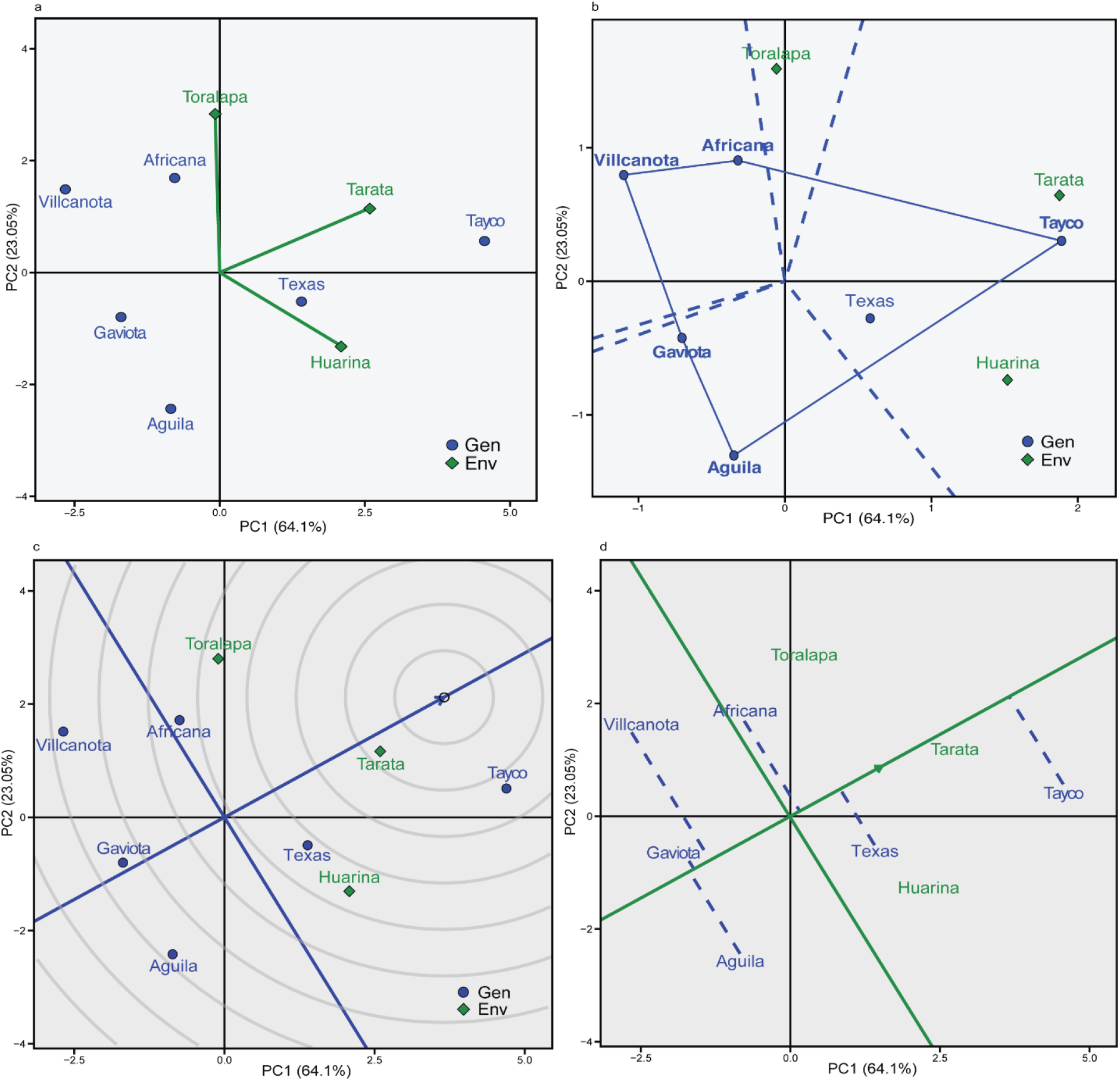
GGE biplot analysis: a) basic biplot, b) who-won-where, c) genotype ranking (ideal genotype) and d) yield vs. genotype stability evaluated in three locations during the 2018-2019 agricultural campaign.

The center of the concentric circles (Figure 1c), represents the position of an ideal genotype, although this statement cannot exist in reality. However, it allows inferences and can be used as a reference for the evaluation of the ideal genotype (Yan & Tinker, 2006). As a result, Tayco was located in one of the concentric circles closer to the ideal genotype. Therefore, it can be considered as an ideal genotype in terms of greater dry matter yield capacity and stability. The Villcanota, Gaviota and Aguila genotypes are far from the ideal genotype and have yields below the general average of the research. On the other hand, in Figure 1d, the Tayco variety had the highest average yield and Aguila the lowest average yield.

The average yields are in the following order: Tayco > Texas > Africana. Africana’s yield was the most variable (less stable), while Tayco and Texas were moderately stable with high dry matter yields.

In AMMI biplot based on the WASSB model (Figure 2a), the cumulative variance of the first two components was 100%, therefore, the results based on mixed models would have a technical and statistical validity of inferences of this research. Also, this value is higher than that obtained by the GGE biplot methodology (Figure 1). Based on these two components, four varieties were identified (Tayco, Aguila, Villcanota, and Africana) winners with a tendency in three environments or four sectors in terms of forage biomass production, which have a similarity of results between both analysis models. On the other hand, Tiago Olivoto, Lúcio, da Silva, Marchioro, et al. (2019) proposes a biplot of four sectors or classes (I, II, III and IV) of genotypes/environments for a joint interpretation of average performance and stability (Figure 2b). The classification in quadrant (I) refers to environments of high discrimination capacity and unstable genotypes of low productivity. Thus, Huarina and Tarata would be environments of high discrimination capacity; and Villcanota classifies as an unstable variety and with a yield below the general average. The Tayco variety is included in section II, according to this analysis, it would be an unstable genotype; however, with biomass productivity exceeding the general average and Toralapa would deserve special attention for its capacity for environmental discrimination. Genotypes in quadrant III could be considered stable due to low WAASB values; however, they are of low biomass productivity. According to Tiago Olivoto, Lúcio, da Silva, Marchioro, et al. (2019), genotypes within the class IV are highly productive and widely adapted; therefore, Texas would be the genotype that meets productivity and stability parameters.

**Figure 2.**
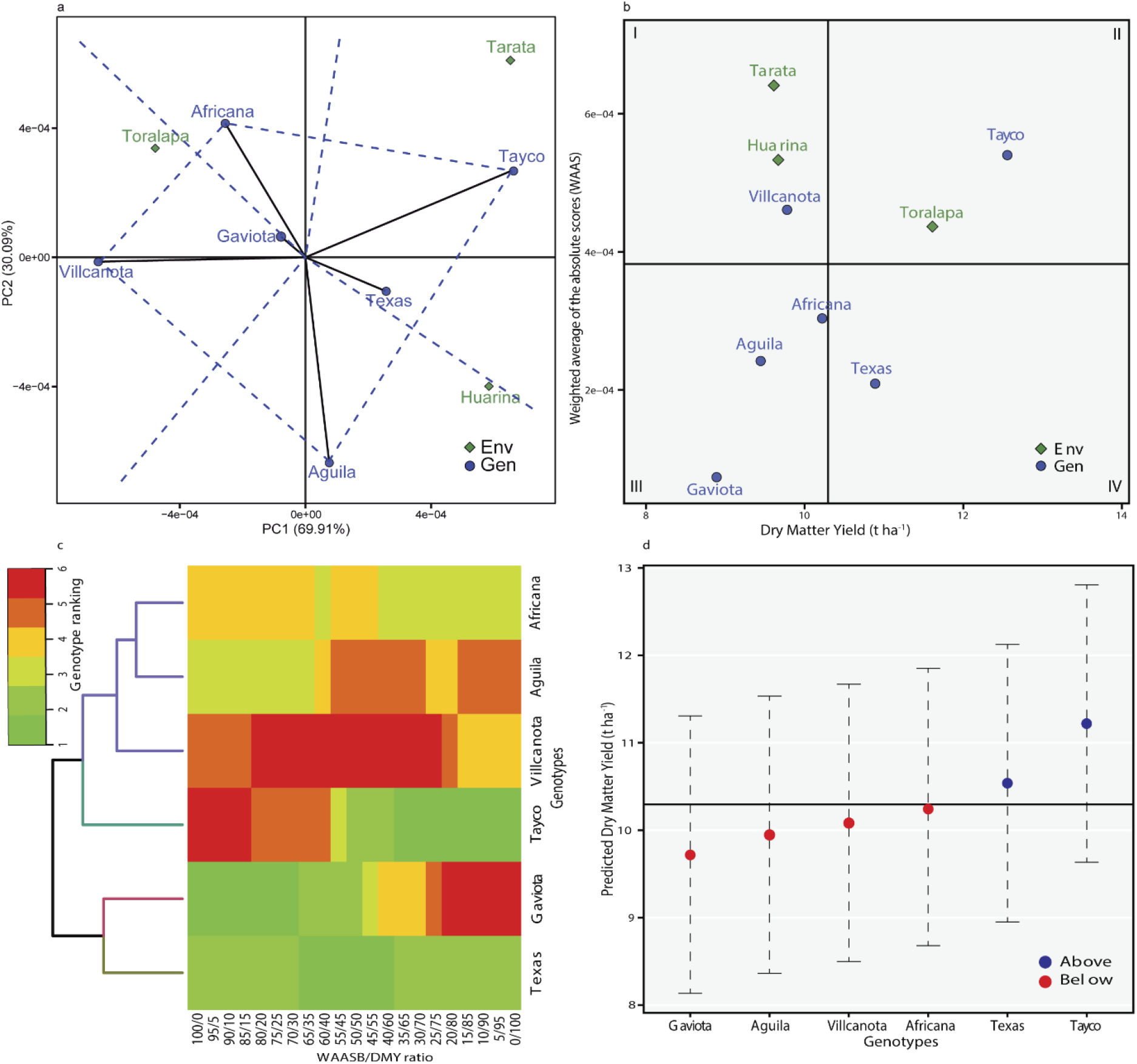
Analysis of multi-environment trials using BLUP (Best Linear Unbiased Prediction): a) AMMI “who-won-where” b) dry matter yield × weighted average of absolute scores (WAASB) c) classification of genotypes according to the weight considered for productivity and stability and d) BLUP plot of six varieties of forage oats evaluated in three locations during the 2018-2019 agricultural campaign.

In Figure 2c referring to the heat map, it shows the genotype classification according to the WAASB/ biomass yield ratio. According to Tiago Olivoto, Lúcio, da Silva, Marchioro, et al. (2019) the ranges obtained with a ratio of 100/0 exclusively consider stability and a ratio of 0/100 considers only the productivity for genotype classification respectively. Thus, the Tayco variety classifies by its characteristic of high productivity and stability, the same trend is observed in figure 2c and 2d of predicted interpretation of BLUP. Also, Texas is listed as a high productivity, which is corroborated by the results in Figure 2c and d.

## Conclusion

Based on two multi-environment analysis methodologies (GGE Biplot and BLUP), two genotypes (Tayco and Texas) were identified with characteristics of high forage productivity and stability. These varieties are recommended to increase forage production in the regions of the valley and the highlands of Bolivia.

